# Cysteine-S-nitrosylation inhibits Rop5-mediated immune evasion in *Toxoplasma gondii*

**DOI:** 10.64898/2025.12.08.693015

**Authors:** SL Lempke, PC DiMare, B Nichols, AM Chaudry, LF Fitzsimmons, ML Reese, SE Ewald

## Abstract

Reactive nitrogen species (RNS) are a mechanism to control microbial infections conserved across the host species of the obligate intracellular parasite *Toxoplasma gondii.* Cysteine S-nitrosylation (SNO) is a reversible post-translational modification that controls complex cell behaviors by regulating protein interactions and signal transduction events. Here we identified a cluster of *T. gondii* secreted effector proteins that are SNO-modified in a host inducible nitric oxide synthetase (iNOS)-dependent manner. Among these were the rhoptry protein 5 (ROP5) paralogs, which are the major virulence determinants in *T. gondii* and an immunodominant antigen in B6 mice. ROP5 was necessary for Type I and Type II parasites to evade IFN-γ-mediated immune clearance in iNOS-deficient macrophages. RNS led to the loss of ROP5 association with the parasitophorous vacuole membrane, which is necessary for the known functions of ROP5. Infection with ROP5 knockout parasites rescued the susceptibility of iNOS-deficient mice to infection with Type II *T. gondii.* Together, these data indicate that RNS can promote cell-autonomous parasite clearance by inhibiting the function of ROP5 paralogs at the parasitophorous vacuole membrane.

**Importance:** RNS are necessary for cell-autonomous immunity to *T. gondii* infection; however, the molecular mechanisms by which RNS regulate parasite control remain poorly understood. Our findings support a model where post-translational modification of ROP5 by RNS is a conserved mechanism of inhibiting the functions of divergent ROP5 paralogs. These data provide a specific example of how host RNS are used to counter *T. gondii* immune evasion effectors that can be applied to understand how nitrosylation regulates the function of other parasite effectors and the role of RNS in the control of other intracellular pathogens.

## Introduction

*Toxoplasma gondii* is an obligate intracellular, protozoan parasite that can infect most nucleated cells (1). The definitive hosts for *T. gondii* are feline species, which shed highly infectious oocysts (1). *T. gondii* infects a wide range of intermediate hosts, including livestock, rodents, and an estimated 30% of the human population, resulting in an infection thought to last the life of the host (2–5). Isolates from Europe and North America are predominantly one of three strains referred to as Type I, -II, and -III (6–9). Type II infection typically results in asymptomatic infection of immunocompetent adults and dose-dependent infection outcomes in inbred strains of mice (10). Type I is hypervirulent in many inbred strains of mice, whereas Type III requires the highest inoculum to induce sickness behavior (11). Virulence is largely determined by parasite effector proteins secreted from the rhoptry, dense granule, and microneme organelles that facilitate the development of the parasitophorous vacuole (PV), nutrient acquisition, and immune evasion.

In intermediate hosts, *T. gondii* infection occurs after ingestion of oocysts or tissue cysts contaminating food or water. *T. gondii* invades the small intestine and converts to rapidly dividing tachyzoites (1, 12, 13). The innate immune response is characterized by NF-κB signaling through activated toll-like receptor (TLR) recognition of *T. gondii* (TLR11, −12), commensal bacteria (TLR2, −4, −7, −9), and or secreted parasite effectors (GRA15, Type II) (14–20). This initiates a Th1 inflammatory cascade characterized by interleukin (IL)-1, IL-12, tumor necrosis factor (TNF), nitric oxide, and interferon-gamma (IFNγ) production (21–27). IFNγ is required for T cell-mediated immunity to *T. gondii* and cell-autonomous control of infection (28–32).

Most murine and human cells can upregulate interferon-stimulated genes that participate in *T. gondii* control (27, 33). Two families of IFNγ-inducible GTPases have been extensively studied in the context of *T. gondii* infection. In mouse cells, the p47 immunity-related GTPases (IRGs) and p65 guanylate-binding proteins (GBPs) oligomerize on the *T. gondii* PV, leading to vesiculation and permeabilization of the organelle (30, 34–37). In human cells, the IRGs are not functionally conserved; however, GBPs are necessary for *T. gondii* control. CIM mice originating from Southeast Asia have divergent alleles of IRGb1-2, which facilitate control of the ‘hypervirulent’ Type I *T. gondii* (38); and several parasite effectors have evolved within species to selectively inhibit IRG proteins, underscoring the importance of this pathway for parasite control.

The parasite pseudokinase Rhoptry 5 (ROP5) is a secreted effector and the major virulence determinant in laboratory strains of mice (7, 9, 39, 40). Virulence-associated alleles of ROP5 interact with the cytosolic face of the parasitophorous vacuole membrane (PVM), and allosterically inhibit IRGa6 oligomerization (41). In addition, IRG-evasion of Type I strains is mediated by ROP5 interactions with virulent alleles of ROP-17, −18, and −39, which phosphorylate the IRG proteins and block IRG oligomerization on the PVM (42–47). Type II parasites encode a cluster of *ROP5* alleles associated with lower virulence, rendering Type II parasites sensitive to successful IRG and GBP attack (38). However, the role of ROP5 in Type II infection is understudied compared to Type I. One report from the Bizik lab found that Type II strains lacking the *Rop5* locus had a small but significant increase in IRGB6 localization to the vacuole but were highly attenuated in vivo (10); however, IRGB6 targeting was not significantly reduced by complementation of a single copy of ROP5A or ROP5C into PruΔ*ku80*Δ*rop5*. In macrophage infection, Type II ROP5 was recently implicated in the phosphorylation of TBK1 and IRF3 leading to polyubiquitination of the innate immune sensor STING when ectopically overexpressed, although the relevance of this mechanism to in vivo infection has not been tested (48).

Until recently, it was speculated that IRG and GBP oligomerization on the PVM were sufficient for cell-autonomous parasite killing in macrophages. However, we recently found that inducible nitric oxide synthase (iNOS) is necessary for efficient clearance of Type II parasites targeted by the GBPs in murine macrophages (49). *Nos2* mRNA can increase by three to four orders of magnitude in mouse macrophage and dendritic cells stimulated with NF-κB drivers and IFNγ (50). NOS enzymes convert L-arginine into nitric oxide (NO) and citrulline (51). NO is a liable signaling molecule that reacts with strong radicals to generate highly reactive RNS. RNS reversibly interrupt cell biological processes, rendering microbes sensitive to immune clearance (51–57). We observed that RNS inhibits *T. gondii* replication within GBP2-positive vacuoles and blocks egress-mediated escape from GBP2-positive vacuoles, but does not alter the percentage of GBP2-positive vacuoles (49). Moreover, PV-localized proteins are labeled with antibodies specific to 3-nitrotyrosine residues, an irreversible post-translational modification induced by RNS (58, 59). Tyrosine-nitration is rare and lacks robust tools for the isolation of modified proteins. In contrast, cysteine-S-nitrosylation (SNO) is a more frequent, reversible post-translational modification, with established pipelines for analysis (60, 61). Although iNOS was identified as a critical mediator of *T. gondii* control in mice in the 1990s, the molecular mechanism by which RNS mediates parasite control is not known (62, 63).

In this study, we leverage tools to stabilize and purify RNS-modified proteins from *T. gondii*-infected macrophages. We identified a cluster of secreted parasite effectors that are post-translationally modified downstream of host iNOS. Among these was ROP5, the major *Toxoplasma* virulence effector (7, 9, 42, 46). Here, we show that SNO occurs on ROP5A, -B and -C alleles. RNS leads to ROP5 dissociation from the parasitophorous vacuole membrane, and results in enhanced clearance of Type II and Type I *T. gondii*. Finally, we show that infection with ROP5-deficient parasites trans-complements the susceptibility of iNOS-deficient mice to Type II *T. gondii* infection. Together, these data indicate that post-translational modification of ROP5 by RNS is an ‘Achilles’ heel’ for parasite immune evasion conserved across ROP5 alleles that facilitates cell-autonomous control of *T. gondii* infection.

## Results

### *T. gondii* secreted effectors are nitrosylated in an iNOS-dependent manner

We hypothesized that RNS-mediated post-translational modifications could promote parasite killing by positively regulating host proteins involved in parasite clearance or by negatively regulating *T. gondii* secreted effectors that mediate immune evasion. To identify host and parasite proteins that are nitrosylated in an iNOS-dependent manner, we used a SNO-Biotin Switch Assay, streptavidin-precipitation, and LC-MS/MS analysis (Figure 1A). To upregulate the transcription of GBPs and iNOS, RAW264.7 macrophage-like cell lines stably expressing Cas9 (WT) were primed with interferon-γ (IFN-γ) and the TLR2 ligand PamCys3K. Uninfected wildtype (WT) RAW, infected WT, and infected iNOS-deficient RAW (*Nos2*^-/-^) samples were harvested at six hours post-infection (Figure 1B-D). The efficiency of the SNO-Biotin switch assay was validated by western blot, where few proteins were eluted from the ‘no-label’ condition (Figure S1A). Nitrosylation is a normal part of homeostatic cell signaling, and a similar number of proteins were isolated in each condition (Figure S1A), however, SNO-proteomes clustered by treatment condition rather than collection date using PCA analysis (Figure 1B). Any proteins identified in the ‘no-label’ condition were excluded from further analysis.

**Figure 1.**
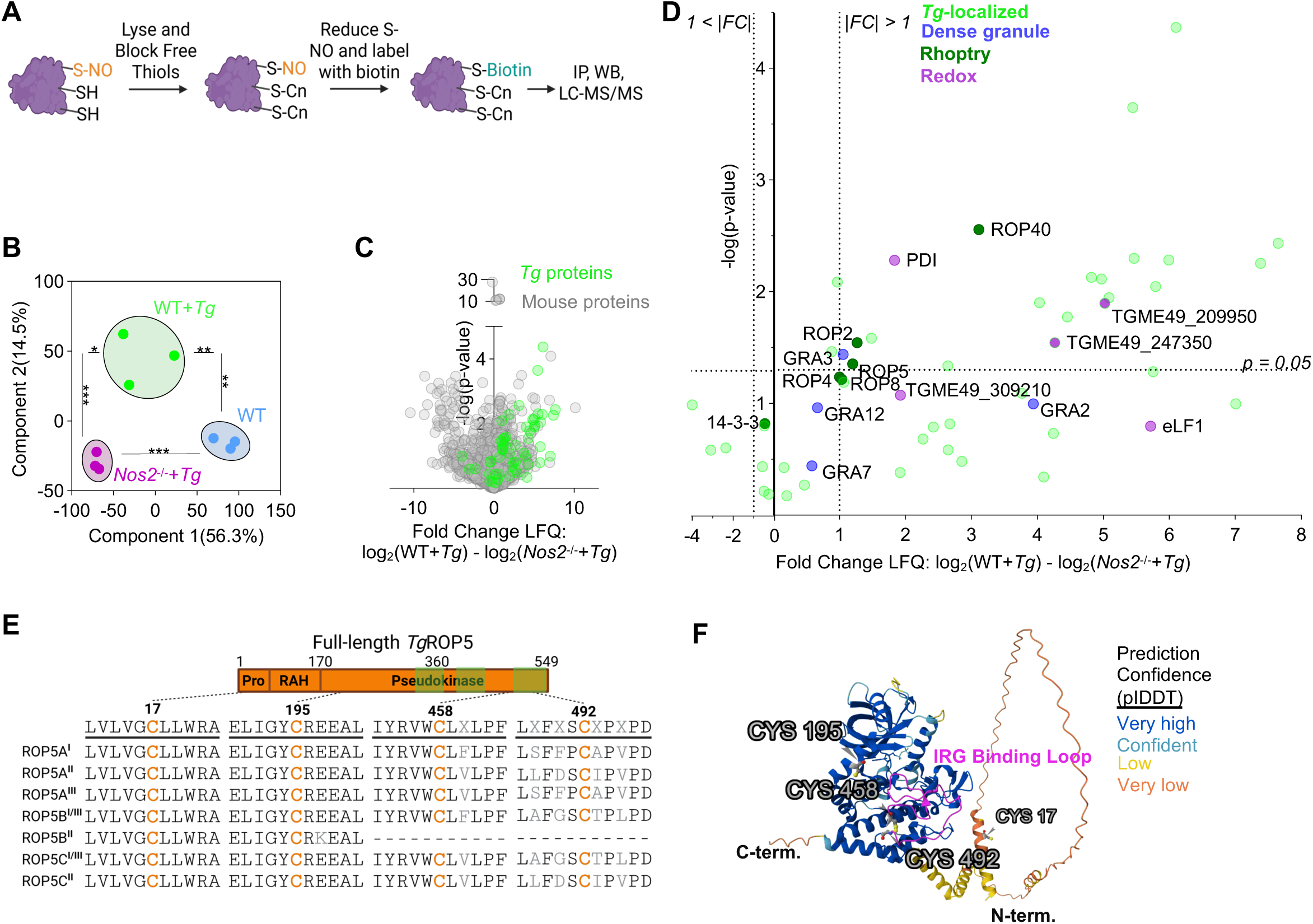
A cluster of *T. gondii* secreted effectors are nitrosylated during infection in an iNOS-dependent manner. **A**) Schematic of SNO-biotin switch assay used to label nitrosylated mouse and *T. gondii (Tg)* proteins 6 hours post infection (hpi) **B-D**) RAW264.7:Cas9 macrophage-like mouse cells (WT) and RAW264.7:Cas9ΔNos2 (*Nos2^-/-^*) were stimulated with 10ng/ml mouse interferon-gamma (IFN-γ) and PAM3CYSK4 24 hours before infection with Type II Me49gLuc at an MOI of 10. Streptavidin-precipitated mouse and *Tg* proteins were identified by LC-MS and label free-quantification. N=3. **B**) Clustering of experimental groups by Principal Component Analysis. Ordinary one-way ANOVA with Tukey’s multiple comparisons test was used on Component 1 (horizontal error bars) and Component 2 (vertical error bars) between groups. **C-D**) Enrichment of S-nitrosylated proteins enriched in infected *Nos2*^-/-^ over infected WT samples. **C**) Average Log2 fold change of mouse (gray) and *Tg* (green) proteins. **D**) Detailed analysis of *Tg* proteins shown in **C**. Labeled proteins are known or predicted *Tg* rhoptry (ROP, dark green), dense granule (GRA, blue) or redox-associated (purple) proteins in data set. Significance analysis represents two-tailed student’s t-test. Inclusion criteria for analysis required a minimum of 1 exclusive unique peptide per protein in two of three replicates, FDR of 0.05 at the peptide and 0.01 at the protein level. *= p<0.05, **= p<0.005, and ***=<0.0005. **E**) Alignment of cysteine (orange) -containing domains of *Tg*ROP5 paralogs expressed in Type I, -II and -III *T. gondii*. Consensus sequence alignment to full-length *Tg*ROP5 with signal-peptide/pro-domain (Pro), arginine-rich amphipathic helical (RAH) domain, and the pseudokinase domain with hypervariable variable regions (green overlay) indicated. *Tg*ROP5B^II^ is truncated at amino acid 344). Dotted lines indicate position of cysteines within *Tg*ROP5. **F**) Position of cysteines in Type II ROP5C structure predicted using AlphaFold 3.

As expected, a subset of SNO-modified host proteins were significantly enriched in infected WT macrophages relative to uninfected (Figure S1B grey, Table S1). This included several proteins with SNO modification sites that have been previously identified experimentally (iNOS(64), RNF213(65), PTPN6(66), PLECTIN(67), IGF2R(68)). We also identified *T. gondii* proteins that were SNO-modified in an iNOS-dependent manner (Figure 1C). In total 54 *T. gondii* proteins were SNO-modified in the data set (Figure S1C, Table S2), the majority of which were more abundant in WT RAW cells relative to iNOS-deficient RAWs (Figure 1C-D, Figure S1C). These included several *T. gondii* proteins predicted to be involved redox signaling (PDI, eLF1, ME49_309210, ME49_209950, ME49_247350) (69). Several proteins were also detected in a SNO proteome study of extracellular, Type I parasites (Figure S1C, *) (70) or are homologs of host proteins regulated by nitrosylation (Figure 1D; Figure S1C, ✝) (ENO2 (71), 14-3-3 (72, 73), HSP/SHP60 (74, 75), LDH (76), RPL18A (77, 78), GAPDH (77, 79, 80), PFKII (70, 81), PDI (70, 82–84), ribosomal proteins).

A cluster of the *T. gondii* proteins that were SNO-modified in an iNOS-dependent manner localize to the *T. gondii* rhoptries or dense granules (Figure 1D, dark green, blue). These proteins are known or predicted to be secreted into the parasitophorous vacuole or host cell, consistent with our hypothesis that nitrosylation promotes *T. gondii* killing by inactivating parasite proteins involved in immune evasion. Also consistent with this hypothesis, many of these secreted effectors were previously found to be dispensable for homeostatic parasite growth in fibroblasts (Figure S1D) (85). Among the SNO-modified, secreted effectors was the pseudokinase ROP5 (Figure 1D).

The *T. gondii Rop5* locus encodes ROP5A, -B, and -C alleles, which arose from paralogous gene expansion (Figure 1E). Type I and Type III parasites express very similar *Rop5A*, *-B,* and *-C* alleles at varying copy numbers (7, 39). In comparison, Type II *Rop5A, -B,* and *-C* alleles are divergent. *Rop5B*^II^ alleles have a premature stop codon before the hypervariable IRG-interaction domain, and Type II parasites encode a higher copy number of *Rop5B^II^*and *Rop5C^II^* alleles than Type I and Type III (7, 9). All *Rop5* alleles, across allelic type and strain, encode 4 conserved cysteines except for *Rop5B*^II^ due to the premature stop codon (Figure 1E-F). In our SNO-precipitation experiment, 5 of the 6 ROP5 peptides identified by LC-MS were conserved across ROP5 alleles, however, one peptide was exclusive to ROP5C^II^ (Table S3). Based on the role of this family as the major determinant of Type I strain hypervirulence in mice (7, 86) and the comparatively understudied role of ROP5 in Type II infection, we next explored whether RNS regulates ROP5 function (10).

### Deletion of the *Rop5* locus rescues the susceptibility of *Nos2*^-/-^ mice to Type II *T. gondii* infection

Mice deficient in iNOS (*Nos2*^-/-^) succumb to Type II *T. gondii* infections that are sub-lethal in wildtype mice (49, 62, 63). We recapitulated this finding using a well-tolerated infection inoculum of 10,000 Pru parasites or a lethal inoculum of 25,000 parasites (Figure 2A). Using this background, we generated a strain lacking the *Rop5* locus (PruΔ*rop5*) (39) and confirmed the Bizik lab study showing that the *Rop5* locus is necessary for Type II virulence in vivo (Figure 2A 2.5×10^4^ Pru vs. Figure 2B 10^7^ PruΔ*Rop5*)(10). Of note, deletion of the *Rop5* locus rescued the impaired immune response of *Nos2^-/-^* mice to infection, as no differences in survival (Figure 2B) or weight loss (Figure 2C) were observed between wildtype B6 and *Nos2^-/-^* mice infected with PruΔ*Rop5* at inoculum that were lethal (10^7^) or primarily sublethal (10^6^) for B6 mice. As shown previously, *Nos2^-/-^* mice had higher Pru parasite load in the lung and spleen 1-week post-infection (Figure 2D). In contrast, the PruΔ*rop5* burdens in the lung and spleen 1-week post-infection), and brain (1-month post-infection) were statistically similar between B6 and *Nos2*^-/-^mice (Figure 2E-F). In contrast, *Nos2^-/-^* mice infected with Pru had a significant increase in parasite load in the lung and spleen at 1-week post-infection (Figure 2D). Taken together, these data are consistent with a model where control of Type II infection in vivo requires iNOS-generated RNS to inactivate ROP5.

**Figure 2.**
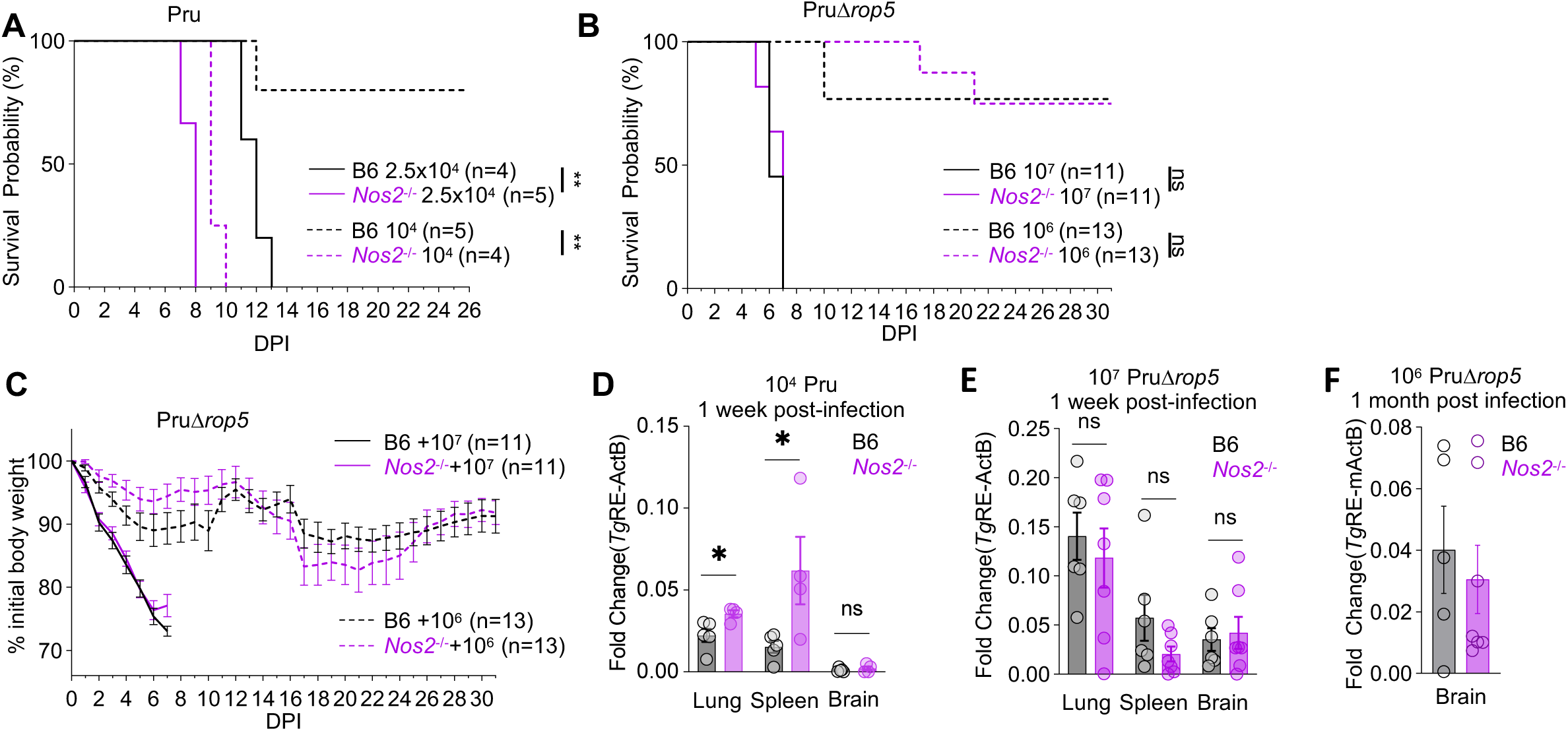
Deletion of the *Tg*ROP5 locus rescues the susceptibility of iNOS-deficient mice *Tg* infection. B6 (WT) and *Nos2*^-/-^ mice were intraperitoneally (i.p.) infected with the parental Pru strain (**A, D**) and PruΔ*rop5* (**B, C, E, F**) and monitored for 31days post-infection or until a humane endpoint was reached. **A**) *Nos2*^-/-^ mice are sensitive to an inoculum of 10^4^ Pru tachyzoites which is non-lethal to B6 mice (B6 n=5, B6 *Nos2*^-/-^ n=4), and more sensitive to an inoculum of 2.5×10^4^ Pru, which is lethal to B6 mice (B6 n=4, n=5). **B-C**) B6 and *Nos2*^-/-^ respond similarly to lethal (10^7^; B6, *Nos2*^-/-^ n=11) and sub-lethal inoculum (10^6^; B6, *Nos2*^-/-^ n=13) of PruΔ*Rop5* in terms of survival (**B**) and weight loss (**C**) (10^6^; B6, *Nos2*^-/-^ n=13). **D**) Pru and **E**) PruΔ*Rop5* abundance (ToxoRE) in tissues relative to mouse was beta-Actin in the Lung, spleen, and brain 6-7 days post infection (10^4^ Pru B6 n=5, *Nos2*^-/-^ n=4; 10^7^ PruΔ*Rop5* B6 n=6, *Nos2*^-/-^ n=7) or **E**) 10^6^ PruΔ*Rop5* at 31 DPI (B6 n=6, *Nos2*^-/-^ n=7). **A-B**) asymmetric survival statistics with Log rank Gehan-Breslow-Wilcoxon test (ns = p>0.05, and **p<0.005) between mouse genotypes within infection dosages. **C-E**) parametric, unpaired t-test (ns p>0.05 and *= p<0.05) between genotypes.

### The *Rop5* locus is necessary for Type II and Type I *T. gondii* to evade cell-autonomous immune clearance by *Nos2*^-/-^ macrophages

The cysteine residues in ROP5 are conserved across strains, so we next asked if there is a role for RNS in the restriction of Type I parasites (RH). As previously reported, by 24 hours post-infection priming with IFNγ or IFNγ and PAM was sufficient to clear over 75% of Type II (Me49-gfp-luciferase) in a manner that was rescued by treatment with the iNOS inhibitor 1400W (Figure 3A, green) (49). In comparison, RH-gfp-luciferase parasites were resistant to killing in IFNγ-primed conditions (Figure 3A, blue), consistent with the ability of RH to partially inhibit host RNS levels (Figure 3B) (87). However, when iNOS was strongly induced by IFNγ and PAM priming (Figure 3B), RH was efficiently cleared in a manner that depended on iNOS activity (Figure 3A, blue).

**Figure 3.**
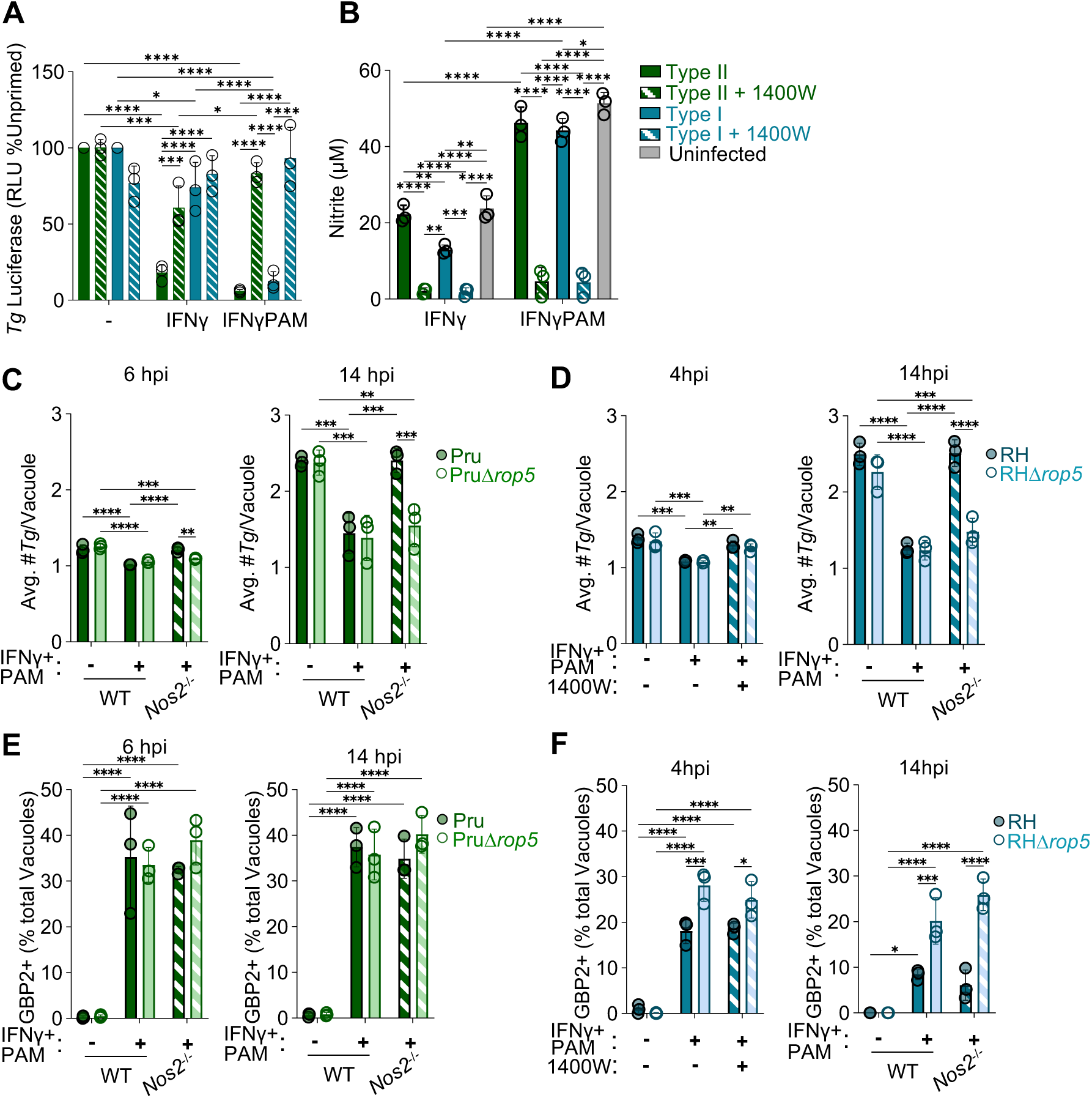
iNOS inhibits Rop5-mediated immune evasion in Type I and Type II *T. gondii*. **A-B**) RAWs were primed with IFNγ with or without PAM3CSK as described in Figure 1 and infected with Me49gLuc and Type I RHgLuc with or without the iNOS inhibitor 1400W (100uM). 24 hours post-infection, parasite abundance was determined by luciferase assay (% of unprimed WT condition within strain) (**A**), or nitrite levels were measured in the supernatants by Greiss assay (**B**). N=3 independent experiments. **C-F**) WT or *Nos2*^-/-^ RAWs were plated on coverslips, primed and infected with Type II Pru and PruΔ*Rop5* (**C, E**) or Type I RH and RHΔ*rop5* (**D, F**). The average number of parasites per vacuole (α-Sag1 staining) was quantified at the indicated time points (**C**-**D**) or the frequency of GBP2-positive vacuoles was quantified (**E-F**). N=3 independent experiments with each replicate containing an average of 379 vacuoles per condition across three fields of view. Ordinary two-way ANOVA with Sidak’s multiple comparison within priming conditions and between strain type (*= p<0.05, **= p<0.005, ***p=<0.0005, and ****p=<0.00005).

We next asked if the iNOS-mediated control was regulated by *T. gondii* ROP5. IFNγ and PAM significantly reduced the frequency of vacuoles containing replicating Pru and PruΔ*rop5* as early as 6 hours post-infection (Figure 3C, left), which was more robust at 14 hours post-infection (Figure 3C, right). In *Nos2*^-/-^ cells, the frequency of vacuoles containing replicating Pru was rescued, and this was dependent on parasite expression of ROP5 (Figure 3C, dashed bars). Similarly, vacuoles containing replicating RH and RHΔ*rop5* parasites were reduced following IFNy and PAMP treatment as early as 4 hours post-infection (Figure 3D, left) and at 14 hours post-infection (Figure 3D, right). In iNOS-deficient RAW cells RH replication was rescued; however, RHΔ*rop5* replication was still controlled in the absence of RNS or iNOS (Figure 3D, dashed bars).

In Type I infection, ROP5 proteins cooperate with Rop17, −18, and −39 to inhibit oligomerization of IRGa6, -b6, and -b10 on the parasitophorous vacuole membrane (42, 46, 47). Several reports have shown that IRG oligomerization is either upstream of, or co-dependent on p65 guanylate binding protein (GBP) recruitment to the vacuole (36, 88, 89). Thus, we evaluated the frequency of GBP2 localization to the vacuole as a proxy for evasion of the IRG/GBP system. GBP2 targeting was similar on Pru and PruΔ*rop5* vacuoles (Figure 3E), even though ROP5-deficient parasites were more susceptible to *Tg* clearance (Figure 3C). There was a small increase in GBP2-positive vacuoles in IFNγ and PAM-primed *Nos2^-/-^*RAWs, however this was not significant (Figure 3E, dashed bars WT vs. *Nos2^-/-^*). This is consistent with our previous report that GBP2-localization is independent of iNOS and is consistent with a minimal role of GBP2 in Type II ROP5 in IRG evasion (10, 38, 44, 49). Consistent with the published role of ROP5, GBP2 localization to RHΔ*rop5* vacuoles was significantly higher than observed on RH vacuoles with IFNγ and PAM-treatment at 4- and 14 hours post-infection (Figure 3F). Similar to our results with Type II, the frequency of GBP2-positive vacuoles was not dependent on iNOS expression (Figure 3F, dashed WT vs. *Nos2^-/-^*). Together, these data indicate that ROP5 is necessary for Type I and Type II to escape cell-autonomous immune clearance in a manner that is inhibited by iNOS.

### Type I ROP5A and Type I/III ROP5B are nitrosylated downstream of host iNOS

To determine if Type I alleles of *Rop5* associated with virulence could be nitrosylated downstream of iNOS, we evaluated RHΔ*rop5* parasites complemented with a *Rop5A* coding sequence from Type I (*Rop5A*^I^-FLAG) and the *Rop5B* allele conserved in Type I and Type III parasites (*Rop5B*^I/III^-FLAG). Both ROP5A^I^ and ROP5B^I/III^ were efficiently precipitated by SNO-biotin switch from infected RAW cells (Figure 4A). ROP5B^I/III^ nitrosylation was significantly reduced when *Nos2*^-/-^ RAWs were infected (Figure 4A-B). These data indicate that RNS may be a mechanism for broadly inactivating ROP5 proteins.

**Figure 4.**
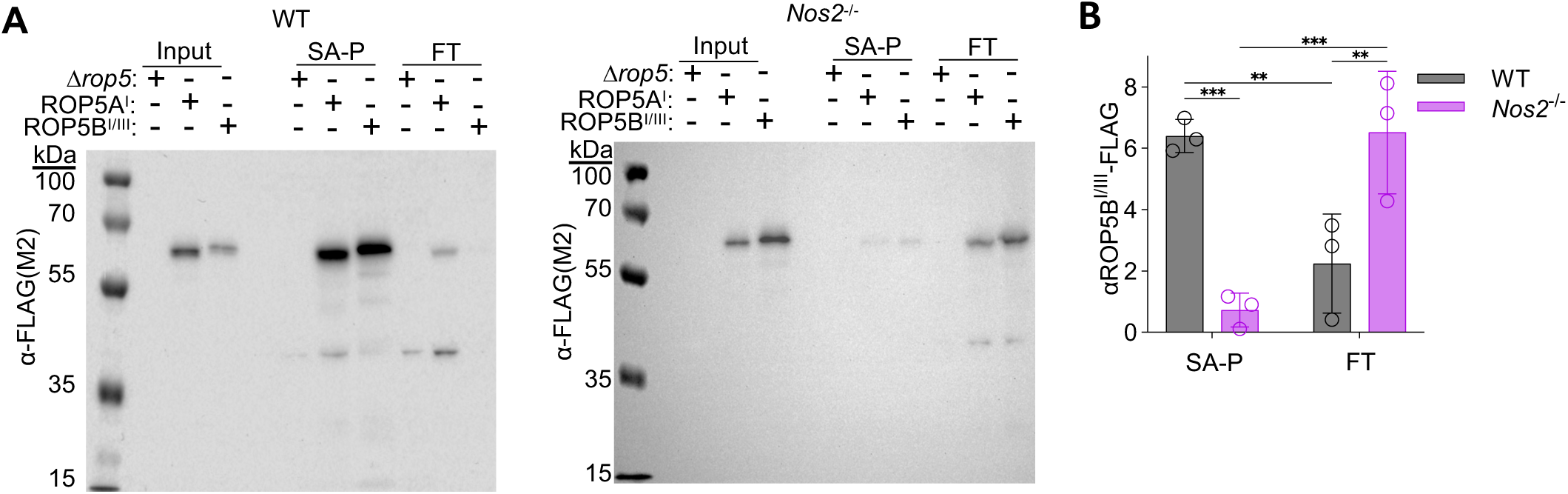
Type I ROP5A and -B alleles are nitrosylated downstream of host iNOS. **A-B**) WT and *Nos2*^-/-^ RAW cells were stimulated and infected with RHΔ*rop5*, RHΔ*rop5*::ROP5A^I^-FLAG (n=2), and RHΔ*rop5:*:ROP5B^I/III^-FLAG (n=3), harvested at 6 hours post-infection, and subjected to SNO-biotin switch assay (Input), biotinylated proteins isolated by Streptavidin Precipitation (SA-P) with collected Flow Through (FT), and western blot as described elsewhere. **B**) Quantification of band intensity for *Tg*ROP5B^I/III^, mouse GAPDH and *Tg*GRA3 shown as the band intensity in the HSS over the total band intensity in HSS and HSP combined for each protein. Ordinary two-way ANOVA with Sidak’s multiple comparison within conditions and host cell type. **= p<0.005 and ***p=<0.0005.

### iNOS inhibits ROP5 association with the parasitophorous vacuole membrane

The known functions of ROP5 depend on interaction with the parasitophorous vacuole membrane (41), so we hypothesized that RNS could interrupt ROP5 membrane association. To assess the sub-cellular localization of ROP5, we infected IFNγ and PAM-primed WT or *Nos2^-/-^* RAW macrophages with RHΔ*rop5::*ROP5B^I/III^-FLAG. Six hours post-infection, samples were syringe lysed and subjected to cellular fractionation. As expected, ROP5 was detected at similar levels in the low-speed pellet of WT and *Nos2^-/-^* RAW cells, which include host nuclei and intact *T. gondii* (Figure 5A, LSP). Similar levels of ROP5 were also detected in the high-speed pellet (HSP) fraction, which contains host and parasite vacuole membrane-associated proteins, including *T. gondii* GRA3 (Figure 5A, HSP). In contrast, ROP5 was identified host-cytosol, high-speed soluble fraction of infected WT, not *Nos2^-/-^* RAWs (Figure 5A-B, mouse GAPDH). The depletion of ROP5 in the cytosolic fraction of *Nos2^-/-^* RAWs was significant compared to WT (Figure 5B), while GAPDH levels were similar. These data are consistent with a model where protein levels of ROP5 in the parasite are not affected by RNS; however, once ROP5 is secreted, RNS causes ROP5 to lose association with the parasitophorous vacuole membrane.

**Figure 5.**
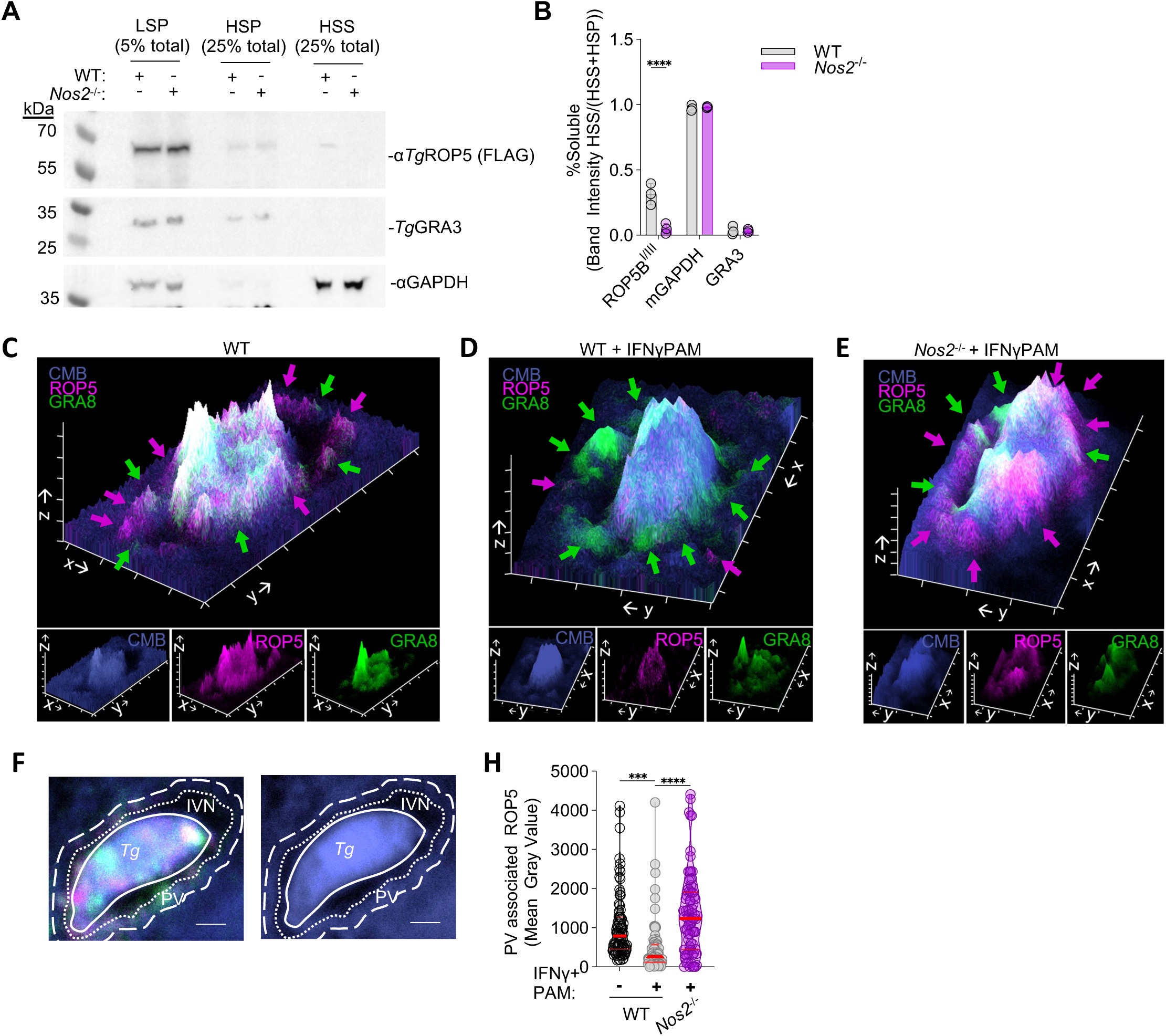
RNS inhibits ROP5 association with the parasitophorous vacuole membranes. **A-B**) WT and *Nos2*^-/-^ RAW macrophages were primed as previously described and infected with RHΔ*rop5*::ROP5B^TypeI/III^-3xFLAG at an MOI of 10. After 4 hours samples were syringe lysed and subject to low-speed centrifugation (LSP, 5% of sample loaded) to eliminate host nuclei and intact parasites. Supernatants were then ultracentrifuged to isolate the membranous, high-speed pellet (HSP, 25% of sample loaded) and the cytosolic, high-speed supernatant was precipitated (HSS, 25% of sample loaded). *T. gondii* ROP5, GRA3 (membrane-bound effector) and host/Tg GAPDH (cytosolic protein) were detected by Western blot (**A**), and the volume-adjusted band densities of each fraction were normalized to the LSP (**B**). **C-H**) WT and *Nos2*^-/-^RAW macrophage were plated on coverslips, primed and infected at an MOI of 2 as described in Figure 3. At 6 hours post-infection samples were fixed and stained with CellMask (Blue), for *T. gondii* GRA8 (green) or ROP5B^I/III^ (anti-FLAG, magenta). Axis tick marks and scale bar = 1µm **C-E**) Representative 2.5D reconstruction of individual parasites from 63xAiryScan images, where each z-stack=0.64nm. Arrows indicate secreted effector localization to the parasitophorous vacuole region. **F-H**) CellMask Blue (CMB) stains the parasite body (Tg; solid white line) and the host cytosol, leaving a CellMask-negative region corresponding to the intravacuolar network region (IVN; between solid and dotted white lines). ROP5B^I/III^ localization to the 0.5 μm region at the edge of the cell mask boundary (‘PV’ between dotted and large dash lines) was quantified (**G**). Each point represents the mean gray value of ROP5B averaged across z-stack of an individual parasite. n= 73-78 vacuoles per condition, collected in three independent experiments. Ordinary one-way ANOVA with Turkey’s multiple comparison test, *** = p<0.005, and ****=p<0.0005.

To test this model directly, we evaluated ROP5B^I/III^ association with the parasitophorous vacuole by immunofluorescence microscopy. CellMask is a reagent that stains the parasite and host cell, resulting in a cell mask-negative region corresponding to the intravacuolar network of the parasitophorous vacuole (37, 49). In unprimed RAW cells, ROP5 localized to the parasite, and the cytosolic face of the cell mask-negative vacuole region (Figure 5C, magenta arrows). The intravacuolar network protein GRA8 partially overlapped with ROP5 localization (Figure 6C, green arrows). IFNγ and PAM-priming significantly reduced the localization of ROP5 to the cytosolic face of the vacuole region but not GRA8 (Figure 5D). ROP5 localization to the periphery of the vacuole was rescued in *Nos2^-/-^* RAW cells primed with IFNγ and PAM (Figure 5E). To quantify ROP5 localization to the region of the parasitophorous vacuole membrane, the mean grey value of the ROP5 signal was quantified in a 0.5 μm region encompassing the cell mask perimeter of each vacuole image in a z-stack (Figure 5F PV, Supplemental Figure 2A-C). This indicated that ROP5 localized to the vacuole perimeter was significantly lower in IFNγ and PAM-primed RAWs compared to *Nos2^-/-^* RAWs and unprimed RAWs (Figure 6H). This loss of ROP5 intensity was also observed when the entire CellMask-negative and vacuole perimeter regions were pooled (Supplementary Figure 2D). By contrast, GRA8 levels were slightly higher in IFNγ and PAM-primed vacuole regions compared *Nos2^-/-^* and unprimed RAWs, indicating that RNS did not trigger a global loss of secreted protein protein association with the vacuole (Supplementary Figure 2E). Together, these data indicate that nitrosylation of ROP5 downstream of iNOS leads to ROP5 dissociation from the PVM, rendering parasites more susceptible to cell-autonomous immune clearance.

## Discussion

The anti-microbial role of reactive nitrogen species is conserved across the intermediate hosts of *T. gondii*. Our data are consistent with the conclusion that RNS can promote parasite killing by inhibiting the function of parasite effectors that mediate immune evasion. Here we show that ROP5 proteins are nitrosylated downstream of host iNOS activity (Figure 1C, 4A-B). Further, ROP5 localization to the PVM, which is essential for the known functions of ROP5, is disrupted following exposure to RNS (Figure 5). These findings are consistent with the better-studied role of RNS in the setting of bacterial pathogens, where RNS reversibly inhibit aspects of bacterial biology that facilitate immune evasion or adaptation to immunological stressors (54–57). This finding does not exclude the possibility that RNS may also positively regulate host effectors that mediate cell-autonomous parasite clearance. In our SNO-proteome study, host candidates for this function may be enriched in infected WT samples compared to uninfected (Supplementary Figure 1B) and *Nos2*^-/-^ samples (Figure 1C). As S-nitrosylation is frequently mediated by transnitrosylation, or the transfer of an NO adduct from one SNO-modified protein to the free thiol of another protein, some of the nitrosylated host proteins may function to selectively modify parasite effectors identified in the study (90).

Our data indicate that SNO may be a mechanism of globally inhibiting ROP5 paralogs based on the conservation of cysteines across ROP5A, -B, and -C alleles and across strains. For Type I alleles of ROP5, our data indicate that RNS either prevents ROP5 association with the PVM or leads to ROP5 release from the PVM (Figure 5), enhancing parasite susceptibility to IRG and GBP targeting (Figure 4) (8, 34, 44, 46). Based on this model, we anticipated that RH-infected *Nos2*^-/-^ RAWs would have less GBP2-targeted vacuoles than WT RAWs, and RHΔ*rop5* infected *Nos2*^-/-^ RAWs would have the highest frequency of GBP2-positive vacuoles (Figure 4F). While there was a trend towards this pattern at 14 hours post-infection, the iNOS-dependent differences were not significant, which may be due to the increased kinetics of IRG/GBP targeting *and* parasite clearance. Live imaging experiments designed to evaluate the frequency of IRG/GBP targeting overtime may resolve this discrepancy; however, we currently do not have *rop5* knock-out parasites on a fluorescent background.

Our data are consistent with a conserved mechanism of ROP5 inactivation by RNS in Type I and Type II infection; however, unlike Type I, Rop5-deficient Type II parasites did not have increased GBP2 localization (Figure 3E). Additionally, our data are consistent with previous reports showing that ROP5A^II^ cannot allosterically inhibit IRGa6(44), and that ROP5B^II^, which has been shown to be the major virulence-determining paralog in Type I parasites(8, 38, 45), is truncated in Type II such that it cannot mediate IRG-inhibition. Although Type II ROP5 knockout parasites were shown to have a small but significant increase in IRGa6 staining (10), it is difficult to reconcile this defect with the magnitude of attenuation in vivo (Figure 2). These data suggest that Type II ROP5 may function via a mechanism independent of IRGs or at least the IRGs and chromosome 3 GBPs that have been studied extensively to date(35, 91). The allelic expansion and evolutionary divergence of ROP5 paralogs within Type II parasites could suggest that these proteins have additional undefined roles in combating in immune evasion. Related, it remains to be determined if RNS inactivates other rhoptry proteins via a similar mechanism to ROP5. For example, ROP2, −4, and −8 were also identified in our SNO proteome (Figure 1D). These proteins are structurally related to ROP5, co-localize to the PVM (9), and have been co-immunoprecipitated with ROP18 (bait), −17, and −5 in Type I infection (46). Together, this suggests that ROP2, −4, and −8 are potential candidates to regulate ROP5 function in Type II parasites, where ROP5 is necessary for *T. gondii* survival in macrophages (Figure 3C), and mice (Figure 2)(10), but appears to function largely independent of IRG(44–46) and GBP evasion (Figure 3E).

One limitation of our study is that we could not identify the nitrosylated peptides using the SNO biotin-switch, streptavidin-precipitation and LC-MS approach. Unfortunately, complementary methodology to stabilize and elute nitrosylated peptides yielded inconsistent results, which is most likely due to the low protein coverage of these proteomics approaches, further confounded by the low abundance of parasite proteins relative to the host proteome (92). Future studies to map the nitrosylation sites by targeted mutagenesis may be informative if the substitution of cysteine to alanine or serine does not result in a fundamental loss of ROP5 function. For example, it is possible that one or both disulfide bridge-forming cysteines are nitrosylated (Figure 1F Cys458, Cys492). Nitrosylation could occur as the protein is secreted in the rhoptries, thereby blocking correct folding; or once ROP5 is exposed to the cytosol leading to disruption of folding and dissociation from the vacuole. While mutagenesis may allow us to confirm that these cysteines are required, misfolding due to disruption of the disulfide bridge would prevent testing if bypassing nitrosylation rescues ROP5 function. Alternatively, a SNO-proteome study on extracellular RH (naturally lysed out of fibroblasts) identified a peptide containing cysteine 195 (Figure 1F), although this most like reflects ROP5 stored in the rhoptries that has been modified by parasite-intrinsic RNS (70).

The importance of RNS in shaping host interactions with *T. gondii* is highlighted by the observation that Type I parasites can inhibit RNS (Figure 3B). Our data recapitulates work from the DaMatta lab, which found that Type I parasites can blunt the synthesis of RNS by 30-50% in RAW and J774 macrophage-like cell lines (93). In this study, *Rop5* deletion led to lower nitrite levels compared to infection with the parental strain, although this was not significant, and RNS levels were not impacted by deletion of parasite effectors that regulate dense granule protein secretion (*asp5* and *myr1*) or inhibit STAT-dependent gene transcription (*ist*) (93). The Knoll lab found that deleting patatin-like phospholipase 1 (*TgPL1)* in Type II parasites increased the frequency of ‘degraded’ vacuoles in macrophages activated with LPS and IFN-γ; this was reversed with NOS-inhibitors or by infecting activated, *Nos2*^-/-^ macrophages (87). PL-1 localizes to the intravacuolar network, suggesting that it may have evolved to counteract the functions of host RNS (87). *T. gondii* has a predicted 58 gene predicted to encode redox-associated proteins (including thioredoxins, glutaredoxins, and protein disulfide isomerases), several of which are necessary for parasite homeostatic functions, and five of which were identified in our SNO proteome (Figure 1D, purple) (69, 94, 95). At least two of the 58 are predicted rhoptry or dense granule proteins (not identified in the SNO-proteome), suggesting that other secreted effectors may mitigate host RNS (69, 96).

The role of host versus parasite redox signaling in the regulation of the other SNO-modified parasite proteins identified in this study remains an open question. Nine of the fifty-four parasite proteins identified localize to the rhoptries or dense granules (Supplementary Figure 1C) and several traffic to the parasite plasma membrane (SAG1, −2A, SRS29C). The majority of the SNO-modified proteins function within the parasite where accessibility to host RNS is comparatively limited by several membranes (parasitophorous vacuole membrane, the intravacuolar network membranes, and parasite plasma membranes) which can block or react with several classes of RNS (53, 97). The high levels of RNS produced by macrophages, coupled with permeabilization of these membranes by the IRG and GBP systems, may facilitate direct modification of proteins sequestered within the parasite by host RNS (33, 34). Additionally, host RNS may enhance parasite redox stress, thereby increasing the modification of these proteins through parasite intrinsic pathways. Notably, 14 of the 54 proteins identified in our study were also detected in a SNO-proteome study of Type I tachyzoites that naturally lysed out of fibroblasts, which may be consistent with the latter scenario for some SNO-proteins (70).

Finally, our in vivo data indicate that iNOS-deficient mice are more susceptible to Type II parasite infection (Figure 2) (49, 62). In addition to its role in parasite evasion of cell-autonomous immunity, discussed above, ROP5 is an immunodominant peptide conferring protective CD8 T cell responses in B6 mice (98). Protective CD8 responses require ERAAP to import and process cytosolic antigens in the endoplasmic reticulum, independent of the Sec22 pathway mediating cross-presentation (99, 100). Infection with parasites expressing a ROP5 protein engineered to be secreted into the host cell cytosol led to enhanced effector CD8 T cell responses in host protection (98). Thus, our data indicate that RNS may promote adaptive immune responses in vivo by increasing ROP5 release into the cytosol (Figure 5) for MHC I presentation to T cells. This may also contribute the susceptibility of *Nos2*^-/-^ mice to infection with WT Type II parasites, in a manner that is mitigated by infection with ROP5-deficient parasites (Figure 2), where alternative epitopes must be used to elicit a protective adaptive immune response.

## Materials & Methods

### Mouse strains and husbandry

C57BL/6 (Jax#000664) and *Nos2^-/-^* (B6.129P2-*Nos2*^tm1Lau^/J, Jax#002609) mice were obtained from Jackson Laboratory (Bar Harbor, ME) and were housed at The University of Virginia vivarium in accordance with University of Virginia Institutional Animal Care and Use Committee, Association for Assessment and Accreditation of Laboratory Animal Care, and Institutional Animal Care (Use Committee Protocol 4107-12-21).

### T. gondii strains

The following strains were used for this study: Me49 expressing GFP and luciferase (Me49-gluc)(101), Type I RH expressing GFP and luciferase (RH-gluc)(101), PruΔ*ku80*Δ*hxgprt* (Pru parental)(102), PruΔ*ku80*Δ*rop5*:HXGPRT (PruΔ*rop5*)(10), RHΔ*ku80*Δ*hxgprt*, (RH parental)(39), RHΔ*ku80*Δ*rop5*:HXGPRT (RHΔ*rop5*)(39), RHΔku80ΔROP5:ROP5(III)A-FLAG:ROP5(III)A-HA (39), and RHΔku80Δ*rop5*:ROP5(III)A-FLAG:ROP5(III)B-HA (39)

PruΔ*ku80*Δ*rop5* were generated as previously described(10). CRISPR-assisted homologous recombination in the PruΔ*ku80*Δ*hxgprt* background using the HXGPRT selection gene flanked with ∼1000 kbp up- and downstream of the *Rop5* locus as previously described(39). Knockouts were selected for with mycophenolic acid and xanthine before confirmation with PCR.

### *T. gondii* in vivo infections

Mice were challenged with the indicated inoculum of Pru or PruΔ*rop5* tachyzoites in 0.2 mL PBS (with Ca^2+^Mg^2+^) by intraperitoneal injection. Before infection, mice were housed on mixed bedding for two weeks to normalize microbiota. Animal weights and clinical scores were assessed daily during acute infections, and on alternating days throughout chronic infections. Briefly clinical score was based on % weight loss, appearance, posture, and activity. Each parameter was scored on a 3 point scale. Animals reaching a 3 in any parameter or a cumulative score of 8 were considered to have reached a humane endpoint and euthanized immediately. The diets of all infected animals were supplemented with soft food when mild dehydration was observed for any animal in the experiment. At 6 or 7 days post-infection, brain, spleen, and lung tissues were harvested from matched B6 and *Nos2^-/-^* mice for genomic DNA extraction. Brains were also harvested at 31 days post-infection to assess chronic parasite burden by PCR. Tissues were homogenized by bead beating using a Qiagen Tissuelyzer for 3 minutes at 25 Hz in 1 uL of UltraPure water/1 mg tissue. Genomic DNA was isolated using DNAeasy Blood and Tissue Kit (Qiagen, 69506). Relative *T. gondii* abundance was measured by quantitative PCR of *T. gondii* 529-bp repeat element (RE) and normalized to mouse beta-actin as previously described (49, 103) using TaqMan primer/probes: 529-bp RE forward, 5′-CACAGAAGGGACAGAAGTCGAA-3′ and reverse, 5′-CAGTCCTGATATCTCTCCTCCAAGA-3′; probe: 5′-CTACAGACGCGATGCC-3 (Integrated DNA Technologies); mouse β-actin: Mm02619580_g (Thermo Fisher Scientific).

### Macrophage cell line generation and maintenance

RAW264.7 macrophages stably expressing RAW264.7:Cas9 macrophage-like mouse cells (WT) and RAW264.7:Cas9ΔNos2 (*Nos2^-/-^*) were generated by lentiviral CRISPR/Cas9 editing as described (49). Cells were cultured in Complete DMEM (Gibco, Waltham, MA) supplemented with 10 mM HEPES (Gibco), 1 mM sodium pyruvate (Gibco), 2 mM L-glutamine (Gibco), 100 U/mL Penicillin, 100 U/mL Streptomycin (Gibco), and 10% heat-inactivated fetal bovine serum (Sigma-Aldrich, St. Louis, MO lot#22M276). Cells were passaged at 37°C and 5% CO_2_ in cell-culture treated tissue dishes. For infections, RAW cells were collected, washed with PBS and plated for experimentation. Cells were allowed to recover overnight before cytokine stimulation.

### Macrophage infections

*T. gondii* tachyozites were passed on confluent human foreskin fibroblasts in Complete DMEM (see above) for up to 12 serial passages. Cell media containing extracellular parasites were discarded. Intracellular *T. gondii* were scraped into fresh media and syringe-lysed by three passages through a 22 g blunt-end needle (Instech Laboratories). *T. gondii* were collected by centrifugation at 310 x g for 8 minutes, resuspended in fresh media, and counted on a hemocytometer.

For parasite load assays, 35,000 cells/well (1.75×10^5^ cells / mL) of WT-Cas9 or RAW *Nos2^-/-^* macrophage were plated in a 96-well plate (Corning 3596). The next day, cells were stimulated with 10 ng/mL mIFNγ (R&D, 485MI100CF) and or 10 ng/mL Pam3CSK4 (Invivogen, tlrl-pms) 24 hours before infection. iNOS was inhibited by 100 µM of 1400W-HCl (Selleck Chemicals, S8337) 1 hour before infection. Cells were infected with an MOI of 5 of Me49-gluc or an MOI of 2 RHgluc. After 24 hours of infection, supernatant nitrite was quantified by Greiss Assay (Thermo Fisher Scientific, G7921) and relative parasite burden measured using Steady-Luc Firefly HTS Assay Kit (Biotium, 30028-L2) according to the manufacturer’s instructions. Chemiluminescence was read with the Cytation 5 plate reader (BioTek, Winooski, VT) and relative light units normalized to unstimulated WT cells for relative *T. gondii* burden.

### Immunofluorescence staining and imaging

WT and *Nos2^-/-^* RAW cells were seeded onto poly-d-lysine (MP Biomedicals)-coated #1.5 thickness, glass coverslips (Harvard Apparatus) at 1.5×10^5^ cells/ml, stimulated, and infected at an MOI:2 as described above. At indicated experimental endpoints, media was aspirated, and samples were fixed with 4% PFA (Electron Microscopy Science) in PBS without Ca^2+^Mg^2+^ (PBS-/-) for 15 minutes at room temperature and stored in PBS at 4°C overnight. Samples were brought to room temperature and permeabilized with 0.1% Triton X-100 (Fisher) in PBS for 30 minutes at room temperature. After one wash with Tris Buffered Saline with 0.1% Tween-20 (TBS-T), samples were blocked with 2% BSA, 2%serum of the host of secondary antibody (Jackson ImmunoResearch Donkey,017-000-001; Goat 005-000-001), and 1:200 TruStain FcX PLUS (Biolegend) in TBS-T for 45 minutes. Coverslips were washed with TBS-T and incubated with primary antibodies for 2.5 hours at room temperature. After 3 washes with TBS-T, fluorescent secondary antibodies were prepared in PBS with 2 μg/ml CellMask Blue (Thermo Fisher Scientific, H32720) and incubated on samples for 1 hour at room temperature. After 3 washes with PBS-/-, coverslips were mounted with Prolong Glass Antifade Mountant (Thermo Scientific, P36984).

Primary antibodies used: α-GBP2 (1:500; Proteintech, 11854-1-AP), α-SAG1 (1:500; Thermo Fisher Scientific, MA518268), α-FLAG(M2) (1:500; Sigma-Aldrich, F1804-200UG), α-GRA8 (1:250; BEI, NR-50269). Secondary antibodies used: Donkey anti-Rabbit IgG(H+L) Highly Cross Absorbed-Alexa Fluor 647 (1:500; Thermo Fisher Scientific, A-31573), Goat anti-Rabbit IgG(H+L) Cross Absorbed-Alexa Fluor 568 (1:500; Thermo Fisher Scientific, A-11011). Coverslips were imaged on the Zeiss LSM 880 (Carl Zeiss) using either 40x (Plan-Apochromat NA1.3, Oil DIC M27) or at 63x (Plan-Apochromat NA1.4, Oil DIC M27) with Airy Scan.

### Immunofluorescence analysis of ROP5 association with the parasitophorous vacuole

Samples were prepared as described above and imaged as z-stack images (0.84 µm) with Airy Scan. After Airy Scan Processing (Zeiss Zen Black) into 16-bit images, individual vacuoles were identified based on excluded CellMask Blue signal from within the vacuole and *T. gondii* was identified by positive CellMask Blue signal. Regions of interest (PV and *T. gondii* body) were drawn manually (ImageJ). ROP5 and GRA8 signal was quantified as mean gray value within the *T. gondii* body, within the PV, and on the PV using a 0.5 µm dilation from drawn PV ROI. Data from z-slices bisecting and individual vacuole were averaged per individual vacuole. Overlapping PVs and extracellular *T. gondii* were excluded from analysis.

### SNO biotin switch assay and streptavidin precipitation

WT and *Nos2^-/-^* RAW cells were seeded at 2.5×10^6^cell/15cm treated tissue culture dish three days before infection. Two days before infection, media was replaced with fresh complete DMEM containing 10% dialyzed, heat-inactivated FBS (Complete DMEM-low Biotin, Gibco) and *T. gondii* strains were passed onto confluent HFFs in Complete DMEM-low biotin. One day prior to infection, host cells were stimulated as described above. Cells were infected with an MOI of 10. At 6 hours post infection, SNO-Biotin Switch (Cayman Chemical) assay was performed in low light, according to the manufacturer’s instructions, with the following modification: the blocking step was extended to 1 hour. The final acetone precipitation was incubated at −30°C overnight.

Streptavidin Precipitation: Biotin-Switched protein pellets were resuspended in RIPA (150 mM NaCl, 5 mM EDTA, 1% NP-40, 1% SDS 50 mM Tris-HCl pH 8) or LC-MS/MS IP Buffer (150mM NaCl, 1%CHAPS, 1 mM EDTA, 5% glycerol) supplemented with 1x protease inhibitor (Sigma-Aldrich, 11836170001) before BCA (Thermo Scientific) protein quantification. Magnetic beads (Thermo Scientific) were washed in low-binding tubes (Thermo Scientific, 3451) with end-over-end mixing for 5 minutes before use with biotin-switched samples. For 50 μL of magnetic beads, 500 μg of SNO-biotin switch proteins were added to a final volume of 500 μL. Streptavidin-precipitation occurred overnight at 4°C with gentle end-over-end mixing. Beads were immobilized on a magnetic stand for 3 minutes. Supernatants were removed and saved as ‘flow through’ controls before beads were gently resuspended by pipetting and washed for 5-10 minutes with end-over-end mixing containing 1x protease inhibitor.

Wash series for western blot wash series: 2x TBS-T(TBS-0.1%TWEEN20), 2x TBS-T+500mM NaCl, 2x TBS-T, and once with H2O+0.5%Tween (H2O). Before the final wash, samples were transferred to new low-protein-binding tubes to limit carryover. For analysis by western blot, proteins were gently eluted with 100 μL 0.1 M Glycine pH 2 for 20 minutes at room temperature with end-over-end mixing. Eluted proteins were neutralized with 15 μL of 1M Tris pH 8, and prepared for western blot, described below.

Wash series for LC-MS/MS analysis: 2x TBS-C (TBS-0.02%CHAPS), 1x TBS-C+500mM NaCl, 1x TBS+500mM NaCl, 1x TBS, and 1x dH2O. Before he final wash, samples were transferred to new low-protein binding tubes and submitted for analysis.

### SNO-peptide identification by Liquid chromatography and mass spectrometry

Samples were submitted to the University of Virginia Biomolecular Analysis Facility. Peptides were digested off streptavidin beads with trypsin and separated chromatographically with a Thermo 75µM x 15cm C18 Easy Spray column and detected with a Thermo Orbitrap Exploris 480 mass spectrometer system with an Easy Spray ion source. Mass spectra data were analyzed with Thermo Discoverer software (Version 2.5.0.400) and MaxQuantLFQ data were analyzed in Perseus (Version 2.0.3.1) using Student’s t-test (permutation-based FDR) (104–106). Modifications from biotin-switch assay (free thiols +125.1amu and biotin-labeled +523.5amu) were accounted for in the analysis (Cayman Chemical). Additional modifications included methionine oxidation and cysteine carbamindomethyl with a limit of 5 modifications per peptide. Peptide Spectrum Match and protein False Discovery Rates were set to 0.01 and protein identification was dependent of minimum 1 unique peptide aligned to either mouse (UP000000589) or *T. gondii* (UP000001529). For inclusion in our downstream analysis, proteins required a minimum of 1 species exclusive, unique peptide in 2/3 replicates.

### Cell fractionation

WT and *Nos2^-/-^* RAW cells were plated into 15cm treated, tissue culture dishes, stimulated, and infected with an MOI 10 as previously described. Four hours post-infection; infected cells were washed and scraped into cold PBS-/-supplemented with 100 µM EDTA. Samples were pelleted at 1100 x *g*, 4°C for 8 minutes before resuspension in 3 mL PBS with protease inhibitor and syringe lysed as previously described. Low Speed Pellets (LSP) were generated at 2500 x g, 4°C for 10 min and supernatants transferred into 2.5 mL, Open-Top Thickwall Polycarbonate Tubes (Beckman Coulter, 349622). High Speed Pellets (HSP) and High Speed Supernatent (HSS) were separated using the TLA-100.3 Fixed Angle Rotor (Beckman Coulter) in the Optima TLX Ultra Centrifuge (Beckman Coulter) at 100,000 x *g* for 2 h at 4°C. HSS was acetone precipitated at a 1:5 sample to acetone ratio overnight before pelleting at 4,000xg for 10 minutes at 4°C. All samples were resuspended in RIPA buffer and shaken for on ice for 1h. Insoluble debris from lysed LSP samples was removed by centrifugation prior to sample preparation.

### Western Blot

Protein samples (see above) were resuspended and boiled in Lithium Dodecyl Sulfate (LDS) (63 mM Tris-HCl pH 6.8) or Sodium Dodecyl Sulfate (SDS) buffer (10µM DTT, 0.0005% bromophenol blue, 10% glycerol, 2% SDS) at 65°C for 10min. Samples were sonicated in a water bath three times for 30 seconds with 2 minutes on ice in between. After SDS gel electrophoresis, proteins were transferred onto PVDF membranes (Thermo Scientific, 88520) using theTrans-Blot Turbo system (Bio-Rad). Membranes were blocked with 4% biotin-free ECL Prime Blocking Reagent (Cytiva) in TBS-T for 1 hour at room temperature, followed by primary antibody incubation for 2 hours at room temperature or overnight at 4°C. After three, five-minute washes with TBS-T, membranes were incubated with HRP-conjugated secondary antibody for 1 hour followed by three washes in TBS-T. Pierce ECL Western Blotting Substrate (Thermo Scientific) was used to visualize the HRP on the membrane and images were acquired by the signal accumulation method on the ChemiDoc Imager (Bio-Rad). Volumetric band intensities were measured using Image Lab software (Bio-Rad, version 6.1). Antibodies used: α-FLAG(M2) (1:500; Sigma-Aldrich, F1804-200UG), α-GRA3 (1:250; BEI, NR-50269), HRP-conjugated-GAPDH (1:5000; ProteinTech, HRP-60004), Peroxidase-Streptavidin (1:7500; Jackson ImmunoResearch, 016-030-084), Peroxidase AffiniPure Goat Anti-Mouse IgG (1:5,000-10,000; Jackson ImmunoResearch, 115-035-002). Band densitometry was performed in Image Lab (BioRad, Version 6.1)

### ROP5 cysteine modeling

Protein sequences were sourced from ToxoDB and models generated using AlphaFold Server Powered by AlphaFold 3.

### Data processing and statistical analysis software

Data was processed in Microsoft Excel unless otherwise noted, and then plotted in GraphPad Prism 10 for the indicated statistical analysis.

## Acknowledgements

We thank Dr. Nick Sherman at the University of Virginia Biomolecular Analysis Facility for performing LC-MS analysis. This work is supported by NIGMS R35GM138381 (S.E.E.); the Welch Foundation (I-2075-20240404) and the Burroughs Wellcome Foundation (G-1021959) (M.L.R.). Conceptualization: S.L.L., S.E.E.; Research Design: M.L.R; S.L.L., S.E.E.; Data analysis: S.L.L., P.C.D., A.C.; Manuscript Drafting and Editing: S.L.L., S.E.E., L.F.F.; Supervision: M.L.R., S.E.E. The authors declare no competing interests.

---

**Supplemental Figure 1:**
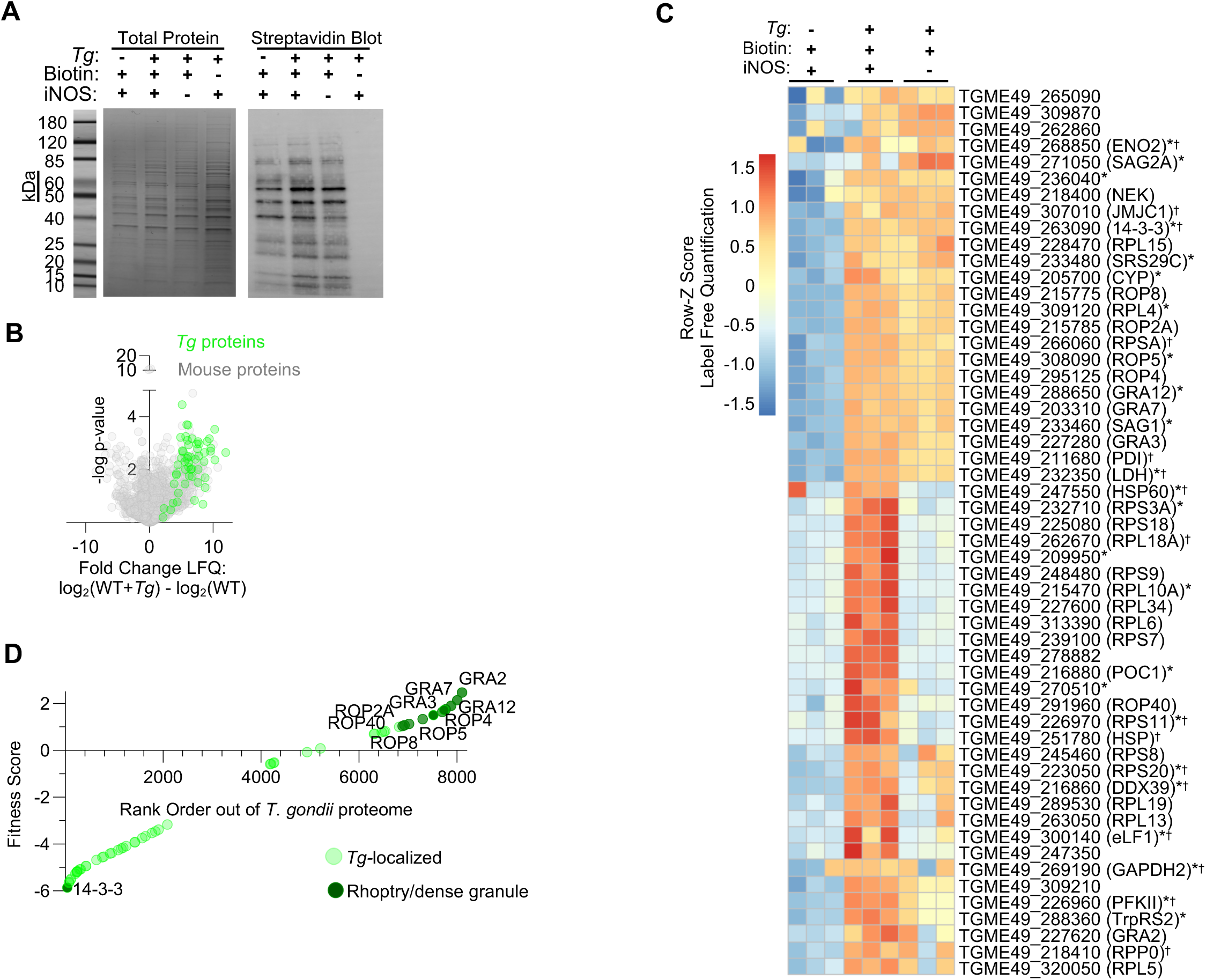
Nitrosylated mouse and *T. gondii* proteins are efficiently labeled and identified by LC-MS. (**A-D**) Nitrosylated mouse and *Tg* proteins were isolated by biotin switch assay as described in Figure 1. **A**) Representative GelCode Blue protein stain (left) and paired streptavidin blot (right) of SNO biotin switch samples isolated for LC-MS. **B**) Average log_2_ fold change enrichment of S-nitrosylated proteins from uninfected WT versus infected WT cells. Mouse proteins (gray) and *Tg* (green) proteins. **C**) All *Tg* proteins identified across three experimental conditions are represented as row-Z score for each protein. * = previously identified as SNO-modified (70) **D**) Nitrosylated *Tg* proteins are ranked based on dispensability during homeostatic growth (fitness score >1 is dispensable (85). Known or predicted *Tg* rhoptry, dense granule, and microneme secreted effectors are indicated in dark green.

**Supplemental Figure 2.**
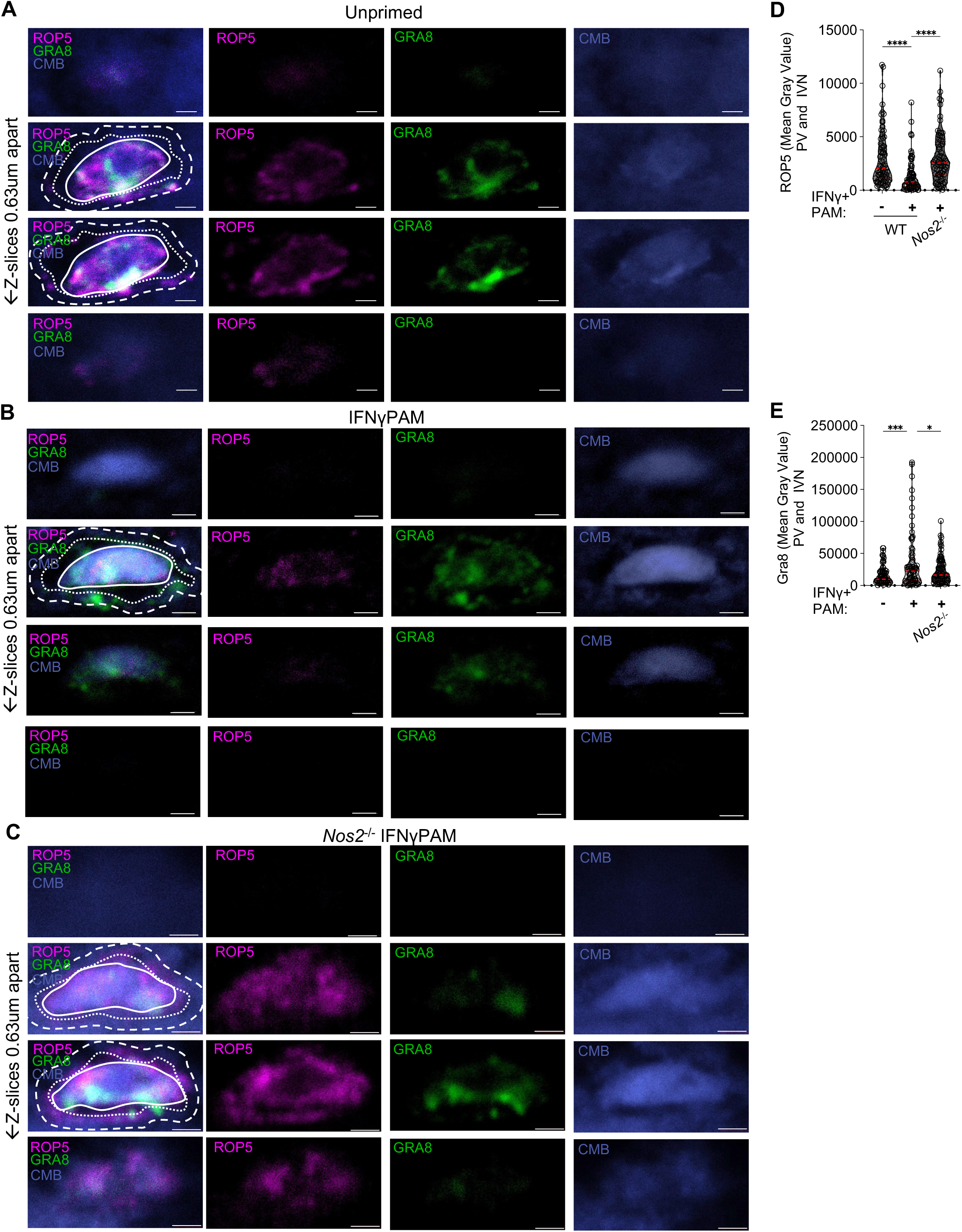
ROP5B localization in representative z-stacks of parasite vacuoles. **A-C)** Individual z-slices from vacuoles shown in Figure 6C-E where *Tg*=solid white line, IVN= region between solid and dotted white lines, and PV-between dotted and large dashed lines. Scale bar = 1 μm. **D-E)** Mean grey value quantification of ROP5B^I/III^ (**D**) or GRA8 (**E**) in the ‘vacuole’ regions pooled between IVN and PV regions. Each point represents the average of z-stack slices of an individual parasite vacuole, N= 73-78 vacuoles per condition, collected from three independent experiments. Ordinary one-way ANOVA with Turkey’s multiple comparison test (ns = p>0.05, *** = p<0.005, and ****=p<0.0005).

## Notes

### Competing Interest Statement

The authors have declared no competing interest.

